# Agent-based modeling of cancer stem cell driven solid tumor growth

**DOI:** 10.1101/035162

**Authors:** Jan Poleszczuk, Paul Macklin, Heiko Enderling

**Affiliations:** Department of Integrated Mathematical Oncology, H. Lee Moffitt Cancer Center & Research Institute, Tampa, FL 33647, USA; Center for Applied Molecular Medicine, University of Southern California, Los Angeles, CA 90033, USA

**Keywords:** agent-based model, tumor growth, cancer stem cell, calibration, domain, search order, high-performance, digital cell

## Abstract

Computational modeling of tumor growth has become an invaluable tool to simulate complex cell-cell interactions and emerging population-level dynamics. Agent-based models are commonly used to describe the behavior and interaction of individual cells in different environments. Behavioral rules can be informed and calibrated by *in vitro* assays, and emerging population-level dynamics may be validated with both *in vitro* and *in vivo* experiments. Here, we describe the design and implementation of a lattice-based agent-based model of cancer stem cell driven tumor growth.

## 1. Introduction

Agent-based modeling has a long history in quantitative oncology ***(1–3)***, including stem cell dynamics ***(4–9)*** and heterogeneity ***(10–12)***. Such models can help predict disease progression and make invaluable recommendations for therapeutic interventions ***(13,14)***. Agent-based models simulate the behavior and interaction of individual cells. Behavioral rules can be dependent on environmental conditions, which includes chemicals in the extracellular environment ***(15–17)***, supporting structures in the extracellular matrix ***(18,19)***, fluid dynamics ***(20)***, physical forces ***(21)***, or presence and interactions with other cells ***(22, 23)***. Computer simulations are usually initialized with a single cell or a cluster of individual cells, and the status and behavior of each cell are typically updated at discrete time points based on their internal rules and current environmental conditions. Such models may help identify if cancer stem cells comprise a subpopulation of specific proportion in a tumor ***(24)***, and how to deliver radiotherapy doses efficiently to eradicate cancer stem cells ***(25)***.

## 2. Materials

Implement all classes and functions in a concurrent version system to allow shared programming and efficient debugging.

### 2.1 Lattice

A finite 2D (or 3D if necessary) lattice, where each site can be occupied by a single or a population of cells (crowding).

1. Lattice size. Simulations are commonly initialized with either *(i)* a small number of cells to observer emergent population-level behavior, or *(ii)* a populated tissue architecture. Define lattice size to accommodate anticipated final cell number. Account for possible boundary effects in case of a growing population. Set the size of a single lattice site to the size of a cell.

2. Neighborhood. Determine if cells interact with their four orthogonal neighbors (north, south, east, west; von Neumann neighborhood), or with the adjacent eight neighboring lattice points (northwest, north, northeast, east, west, southeast, south, southwest; Moore neighborhood) (**Fig. 1**).

**Fig. 1.**
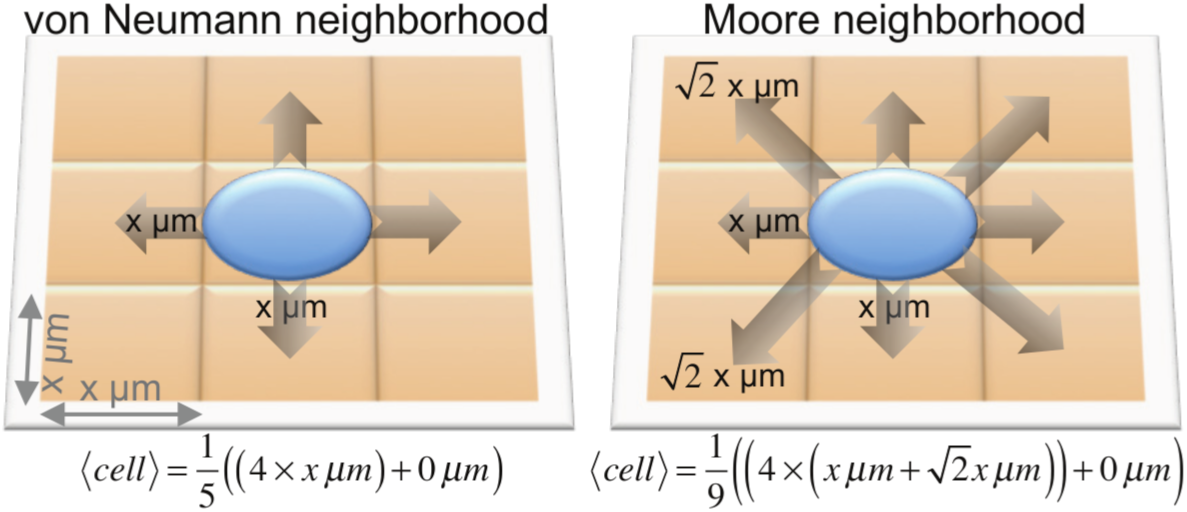
Schematic of expected cell displacement, *〈cell〉*, in the von Neumann (left) and Moore (right) neighborhoods on a two-dimensional lattice with lattice sizes of x^2^ μm^2^.

3. Boundary conditions. Define behavior of cells on the boundary of the lattice, set to either periodic (the boundary “wraps”, so cells on the left edge interact with cells on the far right edge) or no-flux reflective (cells on the edge of the lattice only interact with interior and edge cells). Note that simulations with no-flux boundary conditions may introduce boundary effects, i.e. accumulation of cells near the boundary. For dynamically expanding arrays no boundary conditions are necessary.

### 2.2 Cell

Each cell is an individual entity with the following basic attributes:

1. Time to next division event (t**_c_**).
2. Type of the cell - stem/non-stem (isStem).
3. Probability of symmetric division (p**_s_**). Cancer stem cells divide either symmetrically to produce two identical cancer stem cells, or asymmetrically to produce a cancer stem cell and a non-stem cancer cell ***(26)***. In the case of cancer stem cell define probabilities of symmetric division, 0<p_s_<=1, and asymmetric division p_a_=1-p_s_ (**Fig. 2**).
4. Telomere length (p). Set the telomere length of the initial cell or cell population as a molecular clock ***(27–29)***, which is a quantification of the Hayflick limit ***(30)***.
5. Current cell cycle phase (if required). If information about specific cell cycle phases is required (such as for simulations of cell cycle specific chemotherapeutics) define cell cycle phases. Cell cycle length can be divided into fractions comparable to experimentally measured cell cycle distributions.
6. Probability of spontaneous death (α). Cancer cells (isStem = false) accumulate mutations that introduce genomic instability which may lead to pre-mature cell death. Define a probability of spontaneous cell death, α≥0, at which rate cells may die during cell division attempts. Cancer stem cells (isStem = true) have a superior DNA damage repair machinery ***(31–33)*** and, thus, the probability of spontaneous cell death can be set to zero. For more biological realism, define a cell death rate that is larger than zero, but less than the self-renewal rate to ensure net population growth.

**Fig. 2.**
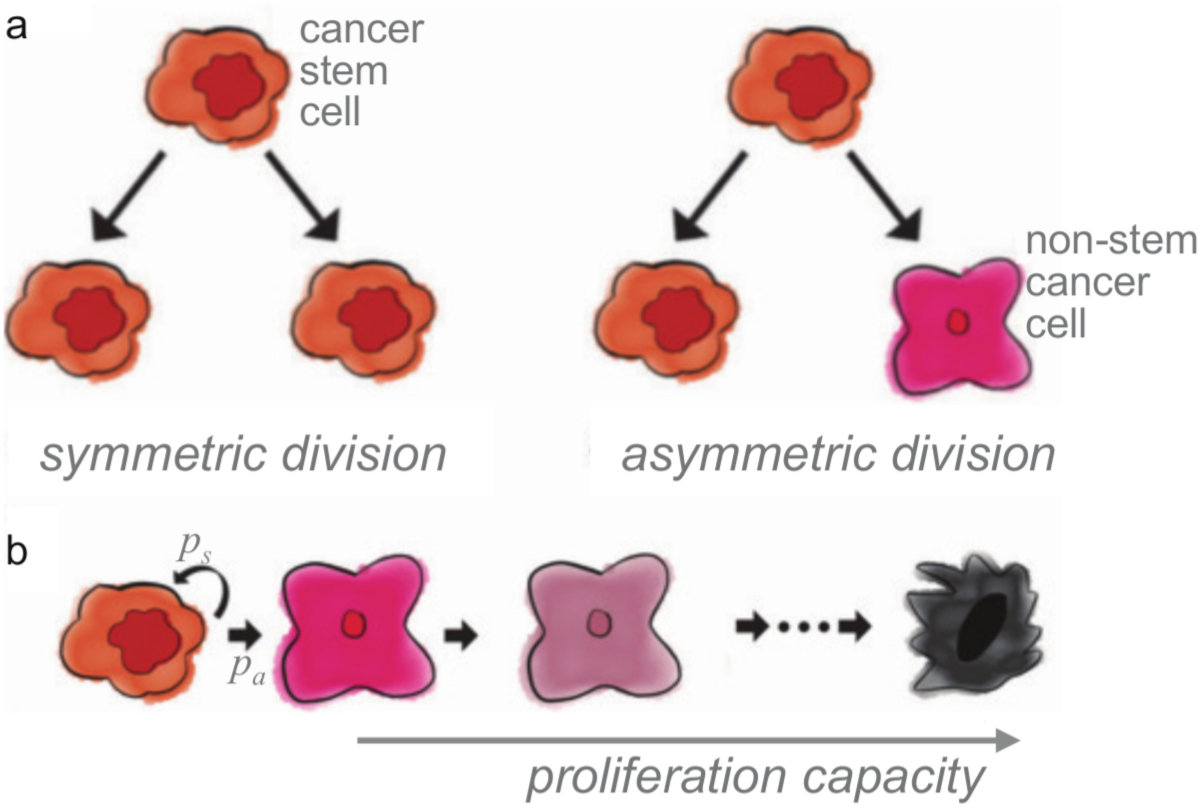
**a)** Schematic of cancer stem cell symmetric and asymmetric division. **b)** Schematic representation of cancer stem cell symmetric *(p_s_*) and asymmetric division *(p_a_*) and non-stem cancer cell proliferation capacity. Adapted from ***(24)***.

Each cell is equipped with a set of basic functions:

1. Procedure *advance time* (input arguments = time increment Δt, list of available sites in the direct neighborhood).

- Decrease time to next division (t_c_) by Δt. If t_c_ < 0 set t_c_ = 0.
- Update current cell cycle phase (if necessary).
- If t_c_ ≤ 0 and there is available space, then perform *division* and *generate new times to next division* for both resulting cells.
2. Procedure *divide* (input argument = list of available sites in the direct neighborhood). Choose a random number 0≤n≤1 from the uniform distribution. If *n* < α, then simulate cell death, i.e. remove cell from the simulation and instantaneously make a corresponding lattice point available. If required, instead of instantaneous cell removal, site on the lattice can be made available for new cells after specified amount of time, e.g. in order to simulate duration of apoptosis process. Otherwise:

- If the cell is not-stem (isStem = false) then decrease the proliferation capacity (p) by one, as non-stem cancer cells do not upregulate telomerase and are thus not long-lived and cannot initiate, retain, and re-initiate tumors ***(34, 35)***. Simulate cell death by removing the cell from the simulation if the proliferation capacity is exhausted, i.e., less or equal to zero. Otherwise, new cell is a clone of the mother cell, i.e. non-stem cancer cell only produces non-stem cancer cell and decremented capacity is inherited by the daughter cell.
- If the cell is cancer stem cell (isStem = true), then draw a number from uniform distribution to decide if the division is symmetric based on the probability p_s_. If division is symmetric, new cell is a perfect clone of the mother cell; that is a cancer stem cells do not erode telomeres and retain identical proliferation capacity after mitosis. Otherwise, non-stem cancer cell offspring of a cancer stem cell inherit the current telomere length and determined proliferation capacity, i.e. new cell is identical except for attribute isStem, which set to false.
3. Procedure *generate time to next division*. Cells divide on average every 24 hours. Derive specific cell cycle times t_c_ (hours), averages and standard deviations from proliferation rate calculations from clonogenic assays or live microscopy imaging ***(36)***.
4. Procedure *random migration* (input argument = list of available sites in the direct neighborhood). Cells may perform a random walk, and probabilities of migrating into adjacent lattice points can be obtained from a discretized diffusion equation ***(37, 38)***. Assuming Moore neighborhood (see section 2.1 above) and a cell at position *(x_0_,y_0_):* at time *t* can move based on available lattice points in the immediate eight-cell Moore neighborhood N(_x0,y0_) with probabilities

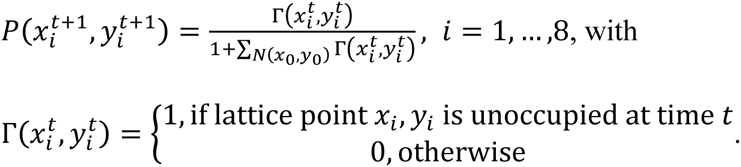 With probability 1/(1+number of unoccupied neighboring lattice points) the cell remains temporarily stationary, and a cell that is completely surrounded by other cells is not moving. Note that in this implementation, movement to all adjacent lattice points is equally weighted. This can be modified to account for increased distance to diagonal lattice points.
5. Procedure *directed migration* (if required) (input arguments = list of available sites in the direct neighborhood, function *F* whose gradient Δ*F* describes the migration stimulus, such as chemoattractant or chemorepellant). Vector 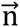 describes the vector connecting the current cell position (*x_0_, y_0_*) to the center of one of the adjacent lattice points *(x_i_,yi)*. Directed migration only occurs towards available lattice sites (S) when 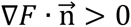. Weight each available site *S* by the factor cos(θ), where θ is the angle between the gradient θ*F* and the movement direction 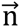 for each possible movement). (Notice that cos(θ) = 1 when Δ*F* and 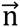 are parallel, and cos(θ) = 0 when they are perpendicular, to give greatest weight to travel along the chemoattractant gradient direction.) Sum each *cos(θ)* over the *S* available sites to get the averaged cosine value *AS*. Define the probability of mobilization *G* through an arbitrary chemotactic/haptotactic responsiveness function via *G* = *H*(|∇*F\AS/(AS +* 1)), where *H* has the properties:

- *H*(0) = 0 (no directed motion when no lattice sites are available or the chemical gradient is zero),
- *H* is a (monotonically) increasing function (directed motility increases with the magnitude chemotactic signal and alignment with the lattice), and
- *H* tends to 1 as |∇*F* |*AS/(AS +* 1) approaches (directed motility increases with the chemotactic signal, but saturates at a maximal level scaled to 1).
6. If cell moves, its direction is weighted by the respective cosines for movement, so that cos(θ)G/(AS) is the probability of movement to the available site at angle θ. This method is explained in detail elsewhere ***(39)***.

## 3. Methods

### 3.1 Programming environment

1. Define programming language. Agent-based models can be implemented and simulated an any programming language; most prominent languages including C++, Java, Julia, Python, Matlab. Each of those languages offers different computational speed and coding feasibility. C++ is considered as the environment offering the best performance, but has a high programming complexity. Matlab offers a great number of built-in functions and is easy to code in, but has significantly slower performance (e.g., when using nested loops).

2. Define graphical output. Visualize simulation solutions of agent-based models using existing implementations of graphical programming or implement specific visualization tools.

3. Agent-based software packages. Utilize pre-developed agent-based software packages; most prominent include Netlogo ***(40, 41)***, CompuCell3D ***(36, 42)***, Chaste ***(43)***, or Swarm ***(44)***.

### 3.3 Simulation procedure

1. Time step. Define the simulation time step, Δt, such that Δt is smaller than the fastest biological process that is being considered in the model. In a model of cell proliferation (~1/day) and cell migration (~1 cell width/hour), set Δt≤1 hour.

2. Develop a simulation flowchart to conceptualize and visualize the simulation procedure. At each defined discrete simulation time step, consider mutually exclusive processes that occur with lowest rate constant first. Let us assume that cell migration and cell proliferation are mutually exclusive (e.g., cells could migrate through most of G1, S, G2 phases). Check if maturation age is reached; if not, check if migration time is reached (**Fig. 3**).

**Fig. 3.**
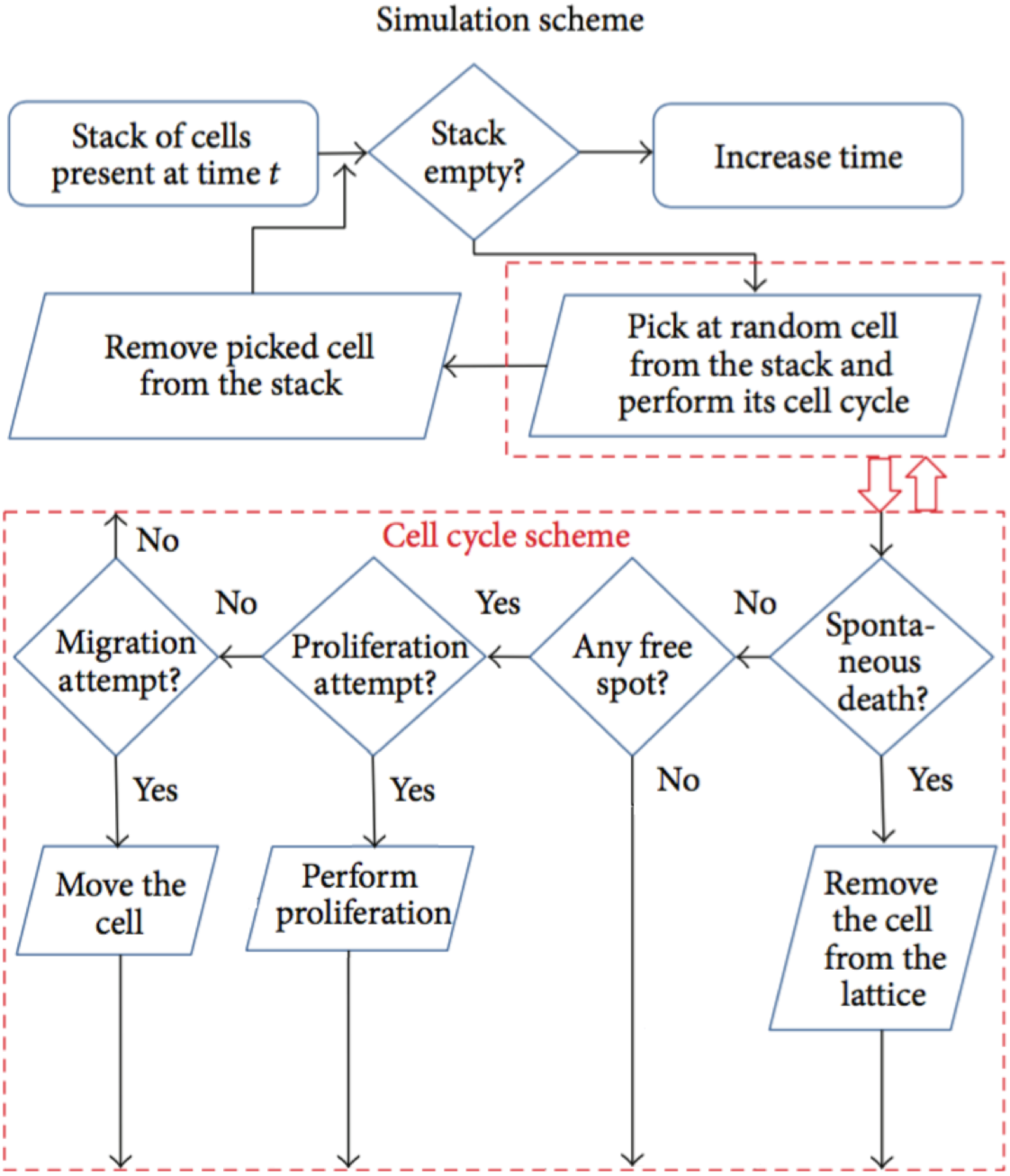
Sample simulation flowchart with the simulation and cell cycle scheme.

3. Lattice update sequence. Update cells in random order to avoid lattice geometry effects. Maintain a list of “live” cell agents and select cells for update from this list at random.

4. Cell update sequence. Consider vacancy in cell neighborhood in random order to avoid lattice geometry effects. Maintain a list of vacant adjacent lattice sites for each cell agent and select for update from this list at random.

## 4. Notes

1. Homogenous populations. The presented model produces a heterogeneous population of stem and non-stem cancer cells. If p_s_=1, the population remains (if only initialized with cancer stem cells) or will become (if initialized with both stem and non-stem cancer cells) homogenous comprised of only cancer stem cells. If p_s_=0, only non-stem cancer cells are produced and the resulting population size oscillates around a dynamic equilibrium or, if the probability of cancer stem cell death is positive and non-zero, the population will inevitably die out ***(45)***.

2. Plasticity. The presented model considers cancer stemness a cell *phenotype*, whereas recent literature may suggest stemness to be a reversible *trait* ***(46, 47)***. The model can be extended to simulated phenotypic plasticity through cancer stem cell differentiation (that is, the cancer stem cell phenotype is set to non-stem cancer cell) and non-stem cancer cell de-differentiation (that is, the non-stem cancer cell phenotype is set to cancer stem cell) ***(48)***.

3. Senescence. The presented model considers cell death after exhaustion of proliferation potential. Alternatively, cells may enter senescence ***(49)*** and, although mitotically inactive, continue to consume resources that influence the behavior of remaining cancer cells ***(50)***.

4. Evolution. The presented model considers fixed parameters or rate constants for each cell trait. Rate constants can change due to mutations and genetic drift ***(51)***, which allows for simulations of the evolution of cancer stem cell traits under different environmental conditions ***(12, 52)***.

5. Carcinogenesis. Such modeling framework can be adapted to model tissue homeostasis and mutation/selection cascades during cancer development ***(16, 53)***.

6. To ensure model results are reproducible, simulation inputs (parameters) and outputs should be recorded using open, standardized data formats, using biologically-driven data elements that can be reused in independent models. The MultiCellDS (multicellular data standard) Project assembled a cross-disciplinary team of biologists, modelers, data scientists, and clinicians to draft a standard for *digital cell lines*, which record phenotype parameters such cell cycle time scales, apoptosis rates, and maximum proliferative capacity. Separate digital cell lines can represent cancer stem cells and non-stem cells. Similarly, standardized simulation outputs can record the position, cell cycle status, and other properties of simulated cells. Using standards will allow better software compatibility, facilitate cross-model validation, and streamline development of user-friendly software to start, modify, visualize, and analyze computational models. See MultiCellDS.org for further details.

7. Model assumptions (e.g., Fig. 3) should also be recorded with standardized data formats. The Cell Behavior Ontology (CBO) ***(54)*** SLUKA provides a “dictionary” (ontology) of applicable stem cell behaviors, but it may not be able to fully capture the entire model logic in its present form. Extensions to the Systems Biology Markup Language (SBML) ***(55)***, such as SBML-Dynamic ***(56)***, are currently being developed to leverage the model structure of SBML and the ontology of CBO to annotate the mathematical structure of agent-based models, but the standards are not yet complete.

8. Treatment. The presented agent-based model can be extended to account for the effects of cancer therapy including radiation ***(57–59)***, chemotherapy ***(60)***, oncolytic viruses ***(61)***, or immunotherapy ***(62)***.

9. Analysis. Agent-based models can be rigorously analyzed for sensitivity and stability ***(63, 64)***. Comparison of coarse-grained model behavior (e.g., growth curves) with known analytical results can quality check calibration protocols and the computational implementation.

10. Dynamically expanding domains. Tradeoffs between lattice size and computing speed can be avoided using dynamically growing domains ***(65)***.

11. Hybrid models. The discussed model setup can be extended to a hybrid discrete-continuous framework where single cells are modeled as discrete agents, and environmental chemicals diffuse on a mapped continuum layer ***(66–68)***. In those models cell cycle time may vary with environmental conditions (e.g., ***(16)*** and ***(69))***.

13. Cell neighborhood. In more complex cases daughter cells can populate empty lattice sites within a given distance to approximate tissue deformability.

## Acknowledgments

JP and HE were partially supported by the Personalized Medicine Award 09-33000-15-03 from the DeBartolo Family Personalized Medicine Institute Pilot Research Awards in Personalized Medicine (PRAPM). PM was supported by the Breast Cancer Research Foundation and the National Institutes of Health [1R01CA180149].

## References

1. A. Anderson, M.A.J. Chaplain, and K. Rejniak (2007) Single-Cell-Based Models in Biology and Medicine, Springer Science & Business Media.

2. H. Enderling and K.A. Rejniak (2013) Simulating cancer: computational models in oncology, Frontiers in oncology. 3, 233.

3. Z. Wang, J.D. Butner, R. Kerketta, et al. (2014) Simulating cancer growth with multiscale agent-based modeling, Seminars in cancer biology.

4. M. d’Inverno and R. Saunders (2004) Agent-Based Modelling of Stem Cell Self-organisation in a Niche, In: Engineering Self-Organising Systems, pp. 52–68 Springer Berlin Heidelberg, Berlin, Heidelberg.

5. D.G. Mallet and L.G. de Pillis (2006) A cellular automata model of tumor-immune system interactions, Journal of theoretical biology. 239, 334–350.

6. D.L. Chao, J.T. Eck, D.E. Brash, et al. (2008) Preneoplastic lesion growth driven by the death of adjacent normal stem cells, Proceedings of the National Academy of Sciences of the United States of America. 105, 15034–15039.

7. H. Enderling, L. Hlatky, and P. Hahnfeldt (2009) Migration rules: tumours are conglomerates of self-metastases, British journal of cancer. 100, 1917–1925.

8. H. Enderling, A.R.A. Anderson, M.A.J. Chaplain, et al. (2009) Paradoxical dependencies of tumor dormancy and progression on basic cell kinetics, Cancer research. 69, 8814–8821.

9. K.-A. Norton and A.S. Popel (2014) An agent-based model of cancer stem cell initiated avascular tumour growth and metastasis: the effect of seeding frequency and location, Journal of The Royal Society Interface. 11, 20140640–20140640.

10. P. Gerlee and A.R.A. Anderson (2008) A hybrid cellular automaton model of clonal evolution in cancer: the emergence of the glycolytic phenotype, Journal of theoretical biology. 250, 705–722.

11. A. Sottoriva, J.J.C. Verhoeff, T. Borovski, et al. (2010) Cancer stem cell tumor model reveals invasive morphology and increased phenotypical heterogeneity, Cancer research. 70, 46–56.

12. J. Poleszczuk, P. Hahnfeldt, and H. Enderling (2015) Evolution and Phenotypic Selection of Cancer Stem Cells, PLoS computational biology. 11, e1004025.

13. A.R.A. Anderson and V. Quaranta (2008) Integrative mathematical oncology, Nature reviews. Cancer. 8, 227–234.

14. T.E. Yankeelov, V. Quaranta, K.J. Evans, et al. (2015) Toward a Science of Tumor Forecasting for Clinical Oncology, Cancer research. 75, 918–923.

15. J.B. Xavier and K.R. Foster (2007) Cooperation and conflict in microbial biofilms, Proceedings of the National Academy of Sciences of the United States of America. 104, 876–881.

16. R.A. Gatenby, K. Smallbone, P.K. Maini, et al. (2007) Cellular adaptations to hypoxia and acidosis during somatic evolution of breast cancer, British journal of cancer. 97, 646–653.

17. H. Enderling, L. Hlatky, and P. Hahnfeldt (2012) The promoting role of a tumour-secreted chemorepellent in self-metastatic tumour progression, Mathematical medicine and biology: a journal of the IMA. 29, 21–29.

18. H. Enderling, N.R. Alexander, E.S. Clark, et al. (2008) Dependence of invadopodia function on collagen fiber spacing and cross-linking: computational modeling and experimental evidence, Biophysical journal. 95, 2203–2218.

19. D.K. Schlüter, I. Ramis-Conde, and M.A.J. Chaplain (2012) Computational modeling of single-cell migration: the leading role of extracellular matrix fibers, Biophysical journal. 103, 1141–1151.

20. K.A. Rejniak (2007) An immersed boundary framework for modelling the growth of individual cells: an application to the early tumour development, Journal of theoretical biology. 247, 186–204.

21. D. Drasdo and S. Höhme (2005) A single-cell-based model of tumor growth in vitro: monolayers and spheroids, Physical biology. 2, 133–147.

22. I. Ramis-Conde, D. Drasdo, A.R.A. Anderson, et al. (2008) Modeling the influence of the E-cadherin-beta-catenin pathway in cancer cell invasion: a multiscale approach, Biophysical journal. 95, 155–165.

23. D.K. Schlüter, I. Ramis-Conde, and M.A.J. Chaplain (2015) Multi-scale modelling of the dynamics of cell colonies: insights into cell-adhesion forces and cancer invasion from in silico simulations, Journal of the Royal Society, Interface / the Royal Society. 12, 20141080–20141080.

24. H. Enderling (2014) Cancer stem cells: small subpopulation or evolving fraction? Integrative biology: quantitative biosciences from nano to macro. 7, 14–23.

25. J.C.L. Alfonso, N. Jagiella, L. Núñez, et al. (2014) Estimating dose painting effects in radiotherapy: a mathematical model, PloS one. 9, e89380.

26. D. Dingli, A. Traulsen, and F. Michor (2007) (A)symmetric stem cell replication and cancer, PLoS computational biology. 3, e53.

27. A.M. Olovnikov (1973) A theory of marginotomy. The incomplete copying of template margin in enzymic synthesis of polynucleotides and biological significance of the phenomenon, Journal of theoretical biology. 41, 181–190.

28. E.H. Blackburn and J.G. Gall (1978) A tandemly repeated sequence at the termini of the extrachromosomal ribosomal RNA genes in Tetrahymena, Journal of molecular biology. 120, 33–53.

29. C.B. Harley (1991) Telomere loss: mitotic clock or genetic time bomb? Mutation ResearchDNAging. 256, 271–282.

30. L. Hayflick (1965) The limited in vitro lifetime of human diploid cell strains, Experimental cell research. 37, 614–636.

31. S. Bao, Q. Wu, R.E. McLendon, et al. (2006) Glioma stem cells promote radioresistance by preferential activation of the DNA damage response, Nature. 444, 756–760.

32. M. Maugeri-Saccà, M. Bartucci, and R. De Maria (2012) DNA damage repair pathways in cancer stem cells, Molecular cancer therapeutics. 11, 1627–1636.

33. S. Skvortsov, P. Debbage, P. Lukas, et al. (2015) Crosstalk between DNA repair and cancer stem cell (CSC) associated intracellular pathways, Seminars in cancer biology. 31, 36–42.

34. R.C. Allsopp, G.B. Morin, R. DePinho, et al. (2003) Telomerase is required to slow telomere shortening and extend replicative lifespan of HSCs during serial transplantation, Hematopoiesis. 102, 517–520.

35. (2010) Telomeres and telomerase in normal and cancer stem cells, 584, 3819–3825.

36. X. Gao, J.T. McDonald, L. Hlatky, et al. (2013) Acute and fractionated irradiation differentially modulate glioma stem cell division kinetics, Cancer research. 73, 1481–1490.

37. A.R.A. Anderson, M.A.J. Chaplain, E.L. Newman, et al. (2000) Mathematical Modelling of Tumour Invasion and Metastasis, Computational and mathematical methods in medicine. 2, 129–154.

38. A.R.A. Anderson, A.M. Weaver, P.T. Cummings, et al. (2006) Tumor morphology and phenotypic evolution driven by selective pressure from the microenvironment, Cell. 127, 905–915.

39. H. Enderling, L. Hlatky, and P. Hahnfeldt (2010) Tumor morphological evolution: directed migration and gain and loss of the self-metastatic phenotype, Biology direct. 5, 23.

40. I. Kareva (2015) Immune evasion through competitive inhibition: the shielding effect of cancer non-stem cells, Journal of theoretical biology. 364, 40–48.

41. R. Bravo and D.E. Axelrod (2013) A calibrated agent-based computer model of stochastic cell dynamics in normal human colon crypts useful for in silico experiments, Theoretical biology & medical modelling. 10, 66.

42. M.H. Swat, G.L. Thomas, J.M. Belmonte, et al. (2012) Multi-scale modeling of tissues using CompuCell3D, Methods in cell biology. 110, 325–366.

43. G.R. Mirams, C.J. Arthurs, M.O. Bernabeu, et al. (2013) Chaste: an open source C++ library for computational physiology and biology, PLoS computational biology. 9, e1002970.

44. N. Minar, R. Burkhart, C. Langton, et al. (1996) The swarm simulation system: A toolkit for building multi-agent simulations, Working Paper 96–06–042, Santa Fe Institute, Santa Fe.

45. H. Enderling (2013) Cancer stem cells and tumor dormancy, Advances in experimental medicine and biology. 734, 55–71.

46. (2013) Cell Plasticity and Heterogeneity in Cancer, 59, 168–179.

47. S. Schwitalla (2014) Tumor cell plasticity: the challenge to catch a moving target, Journal of gastroenterology. 49, 618–627.

48. J. Poleszczuk and H. Enderling (2016) Cancer Stem Cell Plasticity as Tumor Growth Promoter and Catalyst of Population Collapse, Stem cells international. 1–12.

49. J. Campisi, S.H. Kim, C.S. Lim, et al. (2001) Cellular senescence, cancer and aging: the telomere connection, Experimental gerontology. 36, 1619–1637.

50. J. Poleszczuk, P. Hahnfeldt, and H. Enderling (2014) Biphasic modulation of cancer stem cell-driven solid tumour dynamics in response to reactivated replicative senescence, Cell proliferation. 47, 267–276.

51. A. Sottoriva, L. Vermeulen, and S. Tavaré (2011) Modeling evolutionary dynamics of epigenetic mutations in hierarchically organized tumors, PLoS computational biology. 7, e1001132.

52. J.G. Scott, A.B. Hjelmeland, P. Chinnaiyan, et al. (2014) Microenvironmental variables must influence intrinsic phenotypic parameters of cancer stem cells to affect tumourigenicity, PLoS computational biology. 10, e1003433.

53. A.J. Carulli, L.C. Samuelson, and S. Schnell (2014) Unraveling intestinal stem cell behavior with models of crypt dynamics, Integrative biology: quantitative biosciences from nano to macro. 6, 243–257.

54. J.P. Sluka, A. Shirinifard, M. Swat, et al. (2014) The cell behavior ontology: describing the intrinsic biological behaviors of real and model cells seen as active agents, Bioinformatics (Oxford, England). 30, 2367–2374.

55. M. Hucka, A. Finney, H.M. Sauro, et al. (2003) The systems biology markup language (SBML): a medium for representation and exchange of biochemical network models, Bioinformatics (Oxford, England). 19, 524–531.

56. C. Myers and C. Myers (2011) Dynamic Structures in SBML, Nature Precedings.

57. H. Enderling, D. Park, L. Hlatky, et al. (2009) The Importance of Spatial Distribution of Stemness and Proliferation State in Determining Tumor Radioresponse, Mathematical Modelling of Natural Phenomena. 4, 117–133.

58. G.G. Powathil, M. Kohandel, S. Sivaloganathan, et al. (2007) Mathematical modeling of brain tumors: effects of radiotherapy and chemotherapy, Physics in medicine and biology. 52, 3291–3306.

59. H. Kempf, H. Hatzikirou, M. Bleicher, et al. (2013) In silico analysis of cell cycle synchronisation effects in radiotherapy of tumour spheroids, PLoS computational biology. 9, e1003295.

60. G.G. Powathil, K.E. Gordon, L.A. Hill, et al. (2012) Modelling the effects of cell-cycle heterogeneity on the response of a solid tumour to chemotherapy: biological insights from a hybrid multiscale cellular automaton model, Journal of theoretical biology. 308, 1–19.

61. D. Wodarz, A. Hofacre, J.W. Lau, et al. (2012) Complex Spatial Dynamics of Oncolytic Viruses In Vitro: Mathematical and Experimental Approaches, PLoS computational biology. 8,e1002547.

62. W. Shengjun, G. Yunbo, S. Liyan, et al. (2012) Quantitative study of cytotoxic T-lymphocyte immunotherapy for nasopharyngeal carcinoma, Theoretical biology & medical modelling. 9, 6.

63. P. Gerlee and A.R.A. Anderson (2007) Stability analysis of a hybrid cellular automaton model of cell colony growth, Physical review. E, Statistical, nonlinear, and soft matter physics. 75, 051911.

64. P. Gerlee and A.R.A. Anderson (2010) Diffusion-limited tumour growth: simulations and analysis, Mathematical biosciences and engineering: MBE. 7, 385–400.

65. J. Poleszczuk and H. Enderling (2014) A High-Performance Cellular Automaton Model of Tumor Growth with Dynamically Growing Domains, Applied Mathematics. 5, 144–152.

66. A.R.A. Anderson (2005) A hybrid mathematical model of solid tumour invasion: the importance of cell adhesion, Mathematical medicine and biology: a journal of the IMA. 22, 163–186.

67. P. Gerlee and A.R.A. Anderson (2007) An evolutionary hybrid cellular automaton model of solid tumour growth, Journal of theoretical biology. 246, 583–603.

68. H. Enderling, L. Hlatky, and P. Hahnfeldt (2012) Immunoediting: evidence of the multifaceted role of the immune system in self-metastatic tumor growth, Theoretical biology & medical modelling. 9, 31.

69. P. Macklin, M.E. Edgerton, A.M. Thompson, et al. (2012) Patient-calibrated agent-based modelling of ductal carcinoma in situ (DCIS): from microscopic measurements to macroscopic predictions of clinical progression, Journal of theoretical biology. 301, 122–140.

